# Aptamer-Fortified VV-GMCSF-Lact for Systemic Glial Tumor Virotherapy

**DOI:** 10.64898/2025.12.14.694176

**Authors:** Maya Dymova, Anastasia Koshmanova, Natalia Luzan, Evgeny Morozov, Natalya Vasileva, Alisa Ageenko, Victoria Kahanova, Olga Kolovskaya, Tatiana A. Garkusha, Anna Kichkailo, Elena Kuligina, Vladimir Richter

## Abstract

Virotherapy represents a promising approach for cancer treatment. However, when administered intravenously, oncolytic viruses are often hampered by neutralization from antibodies, reducing their antitumor efficacy. Aptamers offer a potential solution by shielding viral particles from neutralizing antibodies and prevent viral particle aggregation. In this study, human glioblastoma xenografts were orthotopically implanted in immunocompromised ICR mice. The therapeutic agents (VV-GMCSF-Lact-Apt1, VV-GMCSF-Lact-Apt2 or VV-GMCSF-Lact) were administered intravenously. Using magnetic resonance imaging, we observed a decrease in tumor growth in all experimental groups. Using the Kaplan-Meier method, it was shown that the mouse survival in the VV-GMCSF-Lact-Apt1, VV-GMCSF-Lact-Apt2 groups was higher than in the control group. The level of key cytokines (IL-1, IL-10, TNF-alpha) in mouse plasma was measured using ELISA. A study of cytokine levels (IL-1, IL-10, and TNF-α) showed that the aptamers shift the immune response from a pro-inflammatory systemic response, by reducing IL-1, to an anti-tumor one, with an increase in TNF-α. In histological analysis, the most destructive changes in tumor tissue were observed in the VV-GMCSF-Lact-Apt2 group, while the morphology of organ tissues (spleen, liver, kidneys and lungs) did not differ between the control and experimental groups. Thus, our results demonstrate a synergistic antitumor effect of the aptamer in combination with the oncolytic virus against glioblastoma tumor xenografts.

## INTRODUCTION

Aptamers are short oligonucleotide sequences of DNA or RNA that exhibit high specificity, selectivity and affinity for target molecules, are easily synthesized and modified ^1^. Characterized by their low molecular weight, low immunogenicity, and low toxicity, these properties underscore their significant potential in both the development of therapeutic agents and diagnostic systems. Nucleic acid aptamers can be rapidly generated through the systematic evolution of ligands using exponential enrichment technology (SELEX), which facilitates efficient binding to a variety of targets, including molecules, viruses, and cells as ligands ^2^. VV-GMCSF-Lact is a promising double recombinant vaccinia virus engineered from the L-IVP strain. It carries deletions in thymidine kinase (tk) and virus growth factor (vgf) genes and insertions of human GMSCF and peptide lactaptin genes into the regions of these deletions, respectively ^3^. These genetic modifications enable the virus to replicate selectively in oncotransformed cells and to induce local immune responses. Phase 1 clinical trials of the recombinant virus VV-GMCSF-Lact as an antitumor agent for breast cancer have been completed (www.clinicaltrials.gov/, NCT05376527). Furthermore, VV-GMCSF-Lact has demonstrated its effectiveness against a broad spectrum of tumor cells of various histogenesis: lung cancer, epidermoid carcinoma, breast cancer and glioblastoma.

The combination of the virotherapy with aptamers holds promise for an enhanced therapeutic effect. Aptamers can shield oncolytic viruses from neutralizing antibodies, act as cryoprotectants in viral suspensions, and reduce viral particle aggregation, thereby increasing their effectiveness ^4^. Aptamers were previously identified that efficiently bound to the VV-GMCSF-Lact virus and demonstrated an synergistic cytotoxic effect against human glioblastoma cells in the presence of serum containing neutralizing antibodies ^5^. Subsequent molecular modeling elucidated the spatial atomic structures of aptamers. They were also shown to reduce the hydrodynamic diameter of viral particles, suggesting decreased interparticle adhesion. The binding efficiency (EC50) of viral particles to these aptamers, along with aptamers cytotoxicity and stability, was thoroughly characterized. Consequently, we selected the most protective aptamers for further study. The preparation VV-GMCSF-Lact-Apt1 was created using aptamer Nv14t_56 (№9 while VV-GMCSF-Lact-Apt2 was generated using the pool of aptamers (NV1t_72 (№1), NV4t_64 (№4), NV4t_53 (№5), NV14t_41 (№8)).

The main objective of this proof-of-principle study was to demonstrate the synergistic effect of aptamers combined with the oncolytic virus VV-GMCSF-Lact in vivo. We used a model with incomplete immunosuppression in mice with orthotopically implanted xenografts. The therapeutic agents (VV-GMCSFLact-Apt1, VV-GMCSFLact-Apt2 or VV-GMCSF-Lact) were administered intravenously. The antitumor efficacy was evaluated by assessing overall survival and analyzing tumor growth inhibition indices via magnetic resonance imaging (MRI). Solid-phase ELISA data on the expression of key cytokines (IL-1, IL-10, TNF-alpha and TGF-β1) were also obtained after the administration of viral preparations VV-GMCSF-Lact-Apt1, VV-GMCSF-Lact-Apt2 and VV-GMCSF-Lact as a control at 24, 72 hours and on the 5th day after administration of the preparation. In histological analysis, the most destructive changes in tumor tissue were observed in the VV-GMCSF-Lact-Apt2 group, while the morphology of organ tissues (spleen, liver, kidneys and lungs) did not differ between the control and experimental groups. In parallel, a histological analysis of tumor tissue in the control and experimental groups and a morphological assessment of organ tissues (spleen, liver, kidneys and lungs) will be performed.

## RESULTS

In this study, we used mice of the ICR line with incomplete immunosuppression. The immunosuppressive regimen consisted of cyclosporine, which inhibits the release of interleukin-1 and interleukin-2—cytokines essential for T-cell activation and proliferation; cyclophosphamide, which depletes neutrophils, B cells, T cells, and natural killer cells; and ketoconazole, an inhibitor of cytochrome P450 that metabolizes cyclosporine. This protocol ensured adequate tumor growth without any histological signs of immune activation for 14 days post-transplantation (Figure 1). A direct effect of cyclophosphamide on the glioma was mitigated because this drug was not administered during the period of extensive tumor growth.

**Figure 1.**
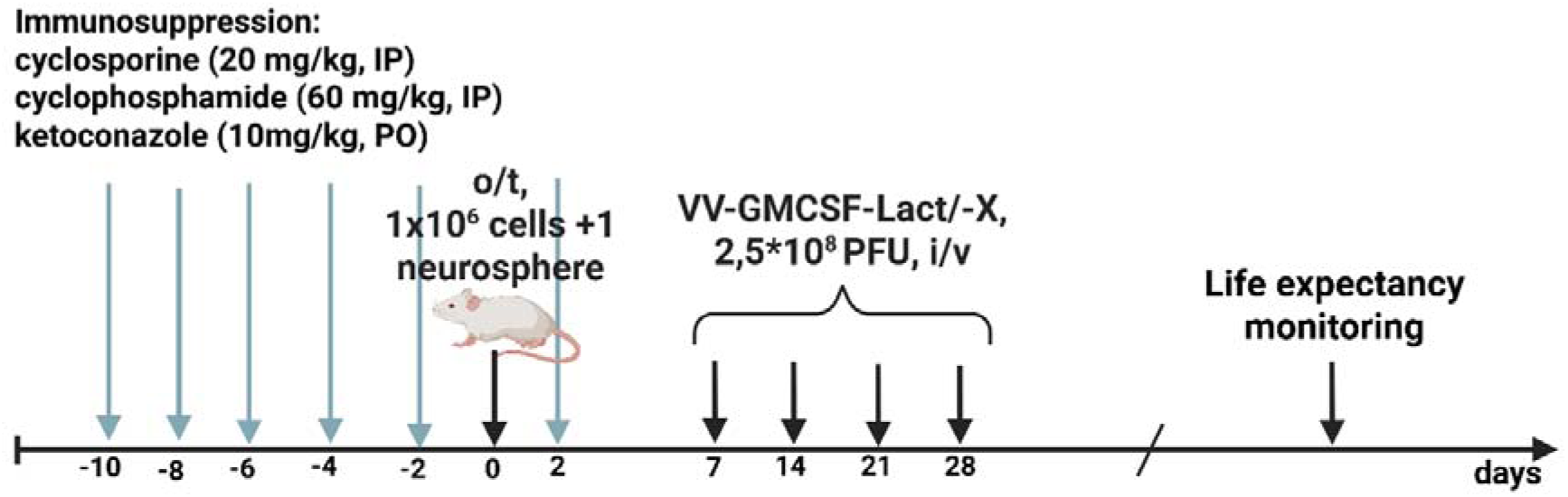
Experimental timeline in mice with incomplete immunosuppression.

Subsequently, primary cell cultures from a human glioblastoma were orthotopically implanted into the right cerebral hemisphere of the mice via stereotactic surgery. The transplantation material consisted of 1×10 adherent cells and one neurosphere in 5 μL of a 1:1 mixture of GrowDex and DMEM medium.

To verify successful xenograft implantation and subsequent growth, magnetic resonance imaging (MRI) with Omniscan contrast agent was performed on two randomly selected animals from each group. Pretreatment MRI scans of all selected animals revealed a zone of heterogeneous contrast enhancement with irregular, poorly defined contours, located cortically and subcortically at the transplantation site. Additional areas of less intense contrast accumulation were visualized in deeper brain regions, potentially corresponding to tumor seeding foci in animals with xenografts. All contrast-enhancing areas were surrounded by zones of vasogenic edema.

### 1. Evaluation of the antitumor efficacy of viral prepararions (VV-GMCSF-Lact, VV-GMCSF-Lact-Apt1, VV-GMCSF-Lact-Apt2) against orthotopically transplanted glioblastoma

#### 1.1. Control group (Saline)

Pre-treatment MRI of control mice receiving intravenous saline revealed contrast-enhancing lesions with irregular borders and perifocal vasogenic edema at the cortical and subcortical transplantation site. Additional areas of contrast uptake in deeper brain regions suggested possible tumor seeding foci (Fig. 2a, b). The initial tumor volumes were 14.1 mm³ (Mouse #1, Fig. 2a) and 6.1 mm³ (Mouse #2, Fig. 2b).

**Figure 2.**
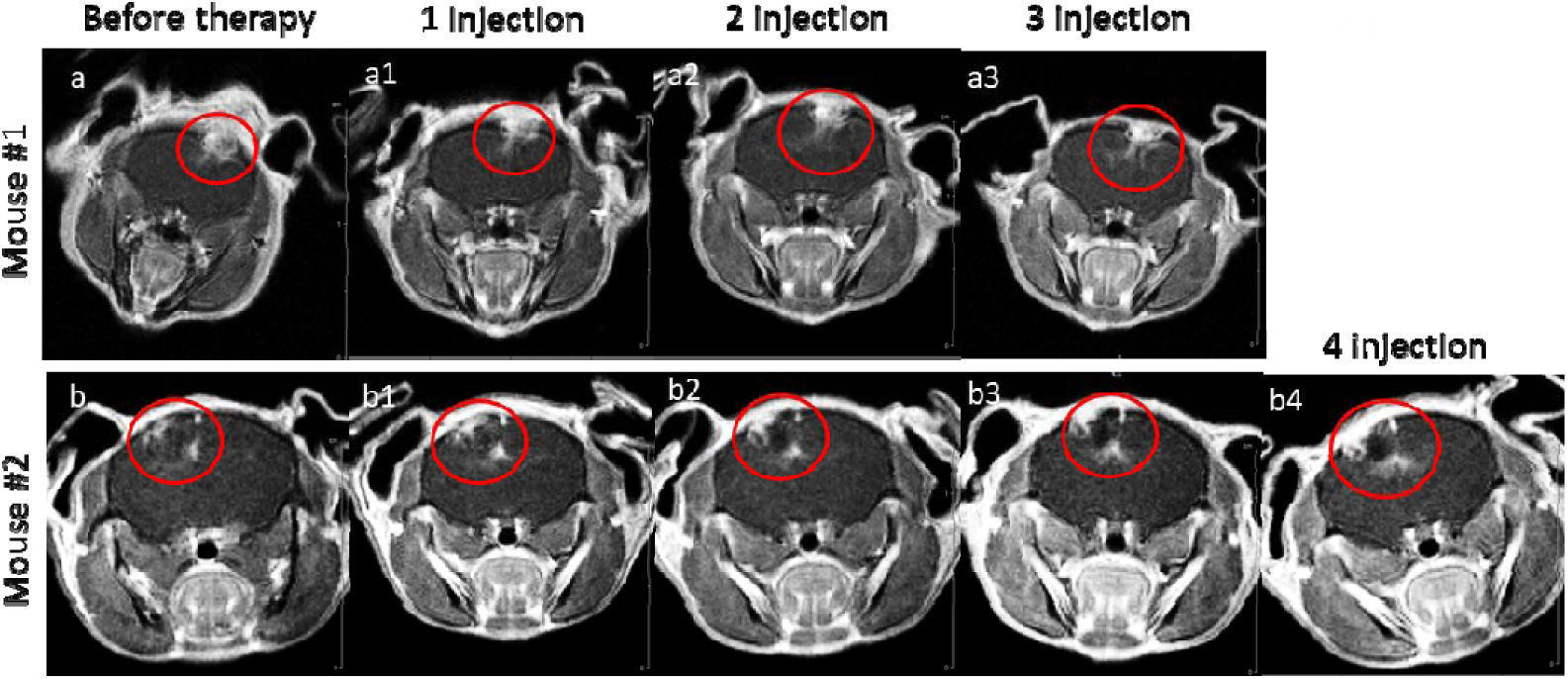
Representative T1-weighted MRI scans of saline-treated control mice: a-a3 – mouse #1, b-b4 – mouse #2; a, b – pre-treatment; a1, b1 – after 1 injection; a2, b2 – after 2 injections; a3, b3 – after 3 injections; b4 – after 4 injections.

One week after saline administration, the contrast-enhancing lesions showed volume expansion (Fig. 2a1, b1), with increased contrast intensity and enlargement of both the main tumor and suspected seeding foci. Tumor volumes increased to 14.9 mm³ (Fig. 2a1) and 6.2 mm³ (Fig. 2b1). After four weeks of saline treatment, further progression was observed with increased lesion volume and perifocal edema, reaching 8.5 mm³ (Fig. 2b4). Animal #1 died before the fourth MRI and 2 months after tumor xenotransplantation. Serial imaging revealed continuous tumor growth, intensified contrast accumulation, and development of cystic-atrophic changes in previously enhanced areas.

#### 1.2. VV-GMCSF-Lact group

Pre-treatment MRI of the VV-GMCSF-Lact group showed intra- and subcortical space-occupying lesions with intense contrast enhancement, perifocal edema, and contralateral seeding foci (Fig. 3a, b). Initial tumor volumes were 26.8 mm³ (Mouse #3, Fig. 3a) and 23.9 mm³ (Mouse #4, Fig. 3b).

**Figure 3.**
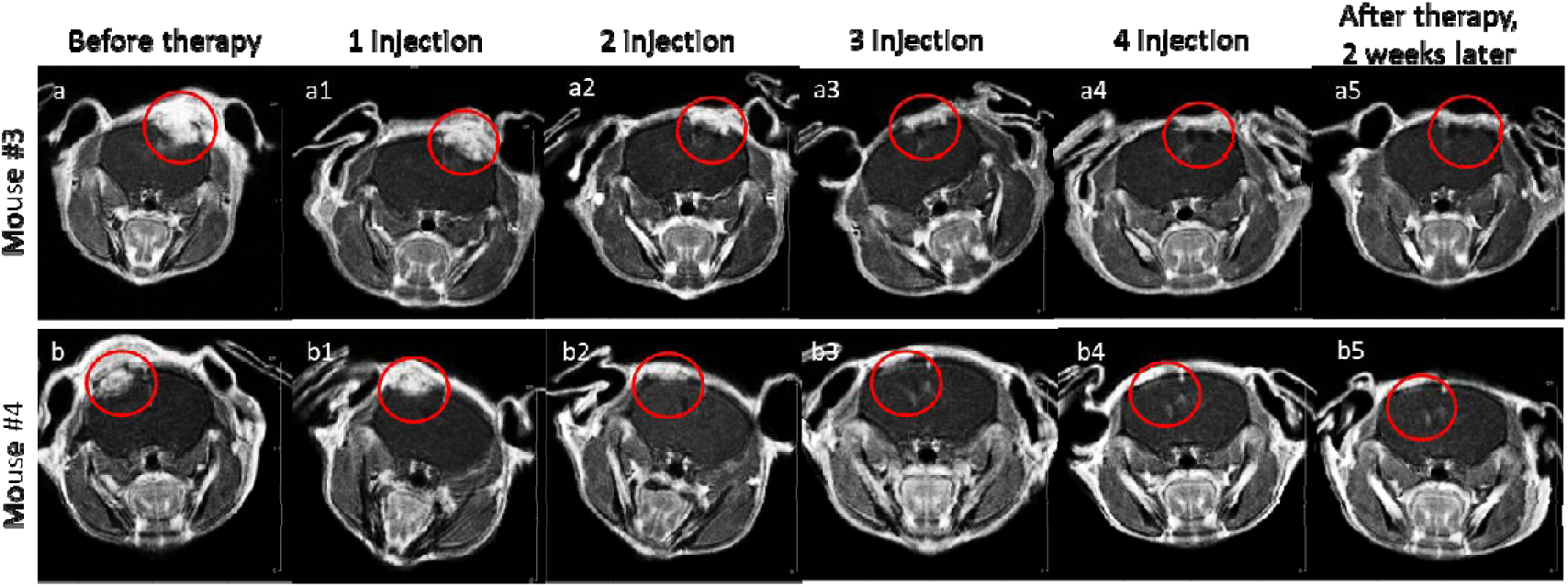
Representative T1-weighted MRI scans of VV-GMCSF-L 564 act treated mice, a course 565 of 4 weekly injections: a-a5 – mouse #3, b-b5 – mouse #4; a, b – pretreatment before therapy; 566 a1, b1 – after 1 injection; a2, b2 – after 2 injections; a3, b3 – after 3 injections; a4, b4 – after 4 567 injections; a5, b5 – 2 weeks after a 4-week treatment course.

After four weeks of therapy, the contrast-enhancing lesions showed significant volume reduction to 9.9 mm³ (63% decrease) and 1.8 mm³ (92.5% decrease), respectively (Fig. 3a4, b4). All sections demonstrated a decrease in the intensity of contrast agent accumulation in the space-occupying formation; in mouse #3, cystic changes formed in the area that had previously accumulated the contrast agent. In mouse #4, a reorganization of the space-occupying process occurred during therapy. The described space-occupying formation after four injections of the oncolytic virus and the accumulation of the contrast agent was practically undetectable. Two weeks after therapy, the formed cystic-atrophic changes persist. Accumulation of the contrast agent and recurrence of the space-occupying process growth were not observed. The volumes of the formations are 9.7 mm3 (Fig. 3a5) and 1.7 mm3 (Fig. 3b5).

**Figure 4.**
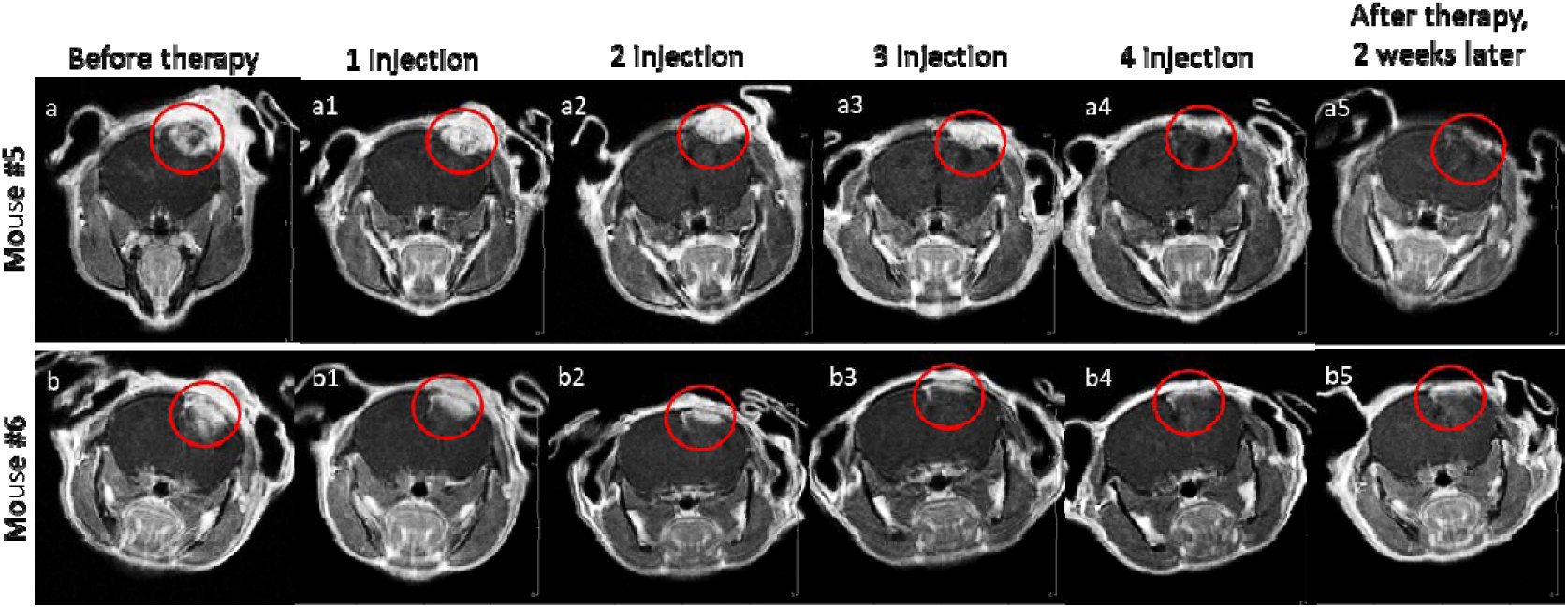
Representative T1-weighted MRI scans of VV-GMCSF-Lact-Apt1 treated mice, a course of 4 weekly injections: a-a5 – mouse #3, b-b5 – mouse #4; a, b – pre-treatment before therapy; a1, b1 – after 1 of treatment; a2, b2 – after 2 weeks of treatment; a3, b3 – after 3 weeks of treatment; a4, b4 – after 4 weeks of treatment; a5, b5 –after 2 weeks following by 4-week treatment course.

#### 1.3. VV-GMCSF-Lact-Apt1 group

Pre-treatment MRI revealed intra- and subcortical lesions with intense contrast enhancement, perifocal edema, pronounced soft tissue components, and deep hemispheric seeding foci (Fig. 4a, b). Initial tumor volumes were 33.7 mm³ (Mouse #5) and 23.2 mm³ (Mouse #6).

After four injections of the viral preparation VV-GMCSF-Lact-Apt1, the previously identified areas of contrast agent accumulation decreased in volume (Fig. 4a4, b4). The volumes of the formations were 11.2 mm3 (Fig. 4a4) and 6.7 mm3 (Fig. 4b4). The decrease in tumor size reached from 22.5 mm3 (67%) to 16.5 (71.1%). Reorganization of space-occupying processes occurs. Space-occupying formations after four injections of VV-GMCSF-Lact-Apt1 significantly decreased. A decrease in the intensity of contrast agent accumulation is visualized, including the soft tissue component of the formation, with the exception of the area of the focus of seeding in the deep sections, cystic changes have formed. The images taken 2 weeks after therapy show no increase in the accumulation of contrast agent in the lesions, the appearance of new areas of contrast agent accumulation, or an increase in the size of the volumetric process. Cystic changes remain in the same volume. The volumes of the formations remained unchanged – 11.0 mm3 (Fig. 4a5) and 6.4 mm3 (Fig. 4b5).

#### 1.4. VV-GMCSF-Lact-Apt2 group

Pre-treatment MRI showed intra- and subcortical lesions with contrast enhancement, perifocal edema, soft tissue components, and deep seeding foci (Fig. 5a, b). Initial volumes were 20.5 mm³ (Mouse #7) and 26.8 mm³ (Mouse #8).

**Figure 5.**
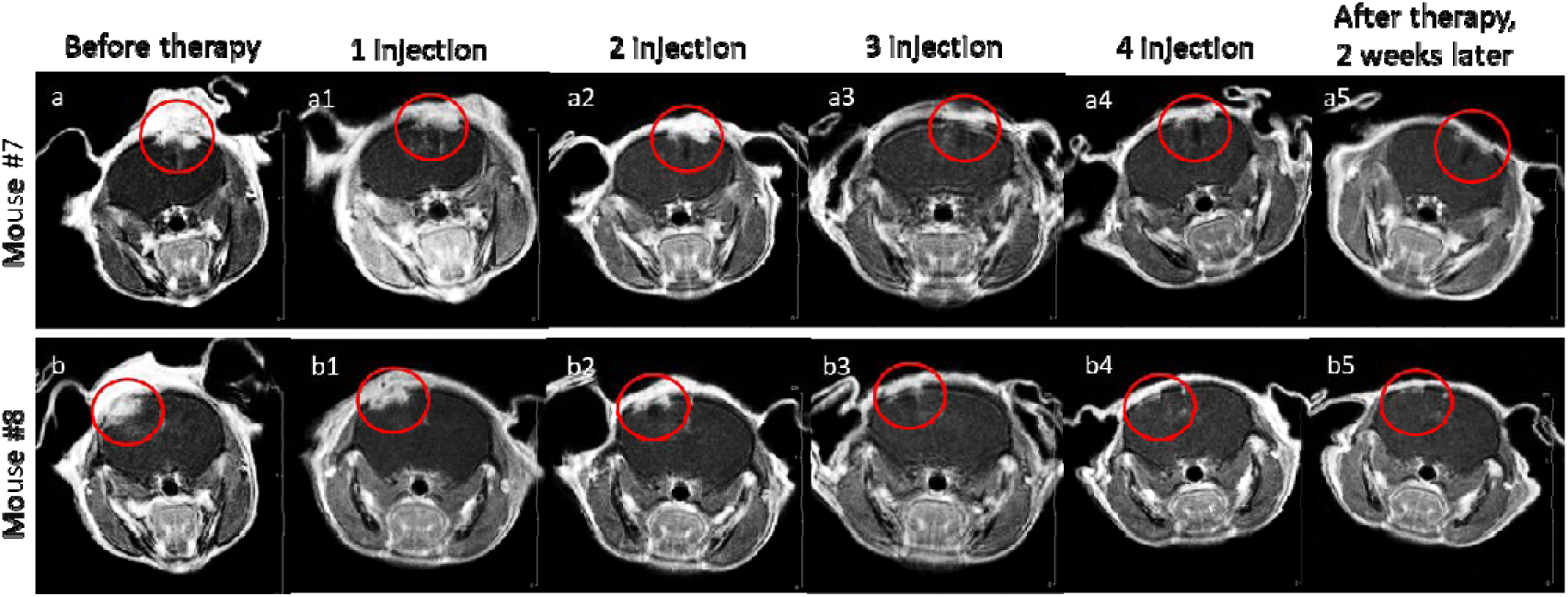
Representative T1-weighted MRI scans of VV-GMCSF-Lact-Apt2 treated mice, a course of 4 weekly injections: a-a5 – mouse #3, b-b5 – mouse #4; a, b – pre-treatment before therapy; a1, b1 – after 1 injection; a2, b2 – after 2 injections; a3, b3 – after 3 injections; a4, b4 – after 4 injections; a5, b5 –after 2 weeks following by 4-week treatment course.

After four weeks of VV-GMCSF-Lact-Apt2 treatment, contrast-enhancing lesions decreased to 5.7 mm³ (72.2% reduction) and 3.4 mm³ (87.5% reduction), respectively (Fig. 5a4, b4). Reorganization of the space-occupying process occurs. The space-occupying formations described before therapy are practically not observed after four injections of the oncolytic virus with a pool of aptamers. A decrease in the intensity of contrast agent accumulation is visualized, including the soft tissue component of the formation, cystic changes have formed. Two weeks after therapy, the formed cystic-atrophic changes persist. Accumulation of contrast agent and relapse of the space-occupying process growth are not observed. The volumes of the formations are 5.3 mm3 (Fig. 5a5) and 3.3 mm3 (Fig. 5b5). Figure 6 shows the dynamics of changes in tumor volumes in different experimental groups during a course of treatment of animals with the viral preparations VV-GMCSFLact-Apt1, VV-GMCSF-Lact-Apt2 and VV-GMCSF-Lact.

**Figure 6.**
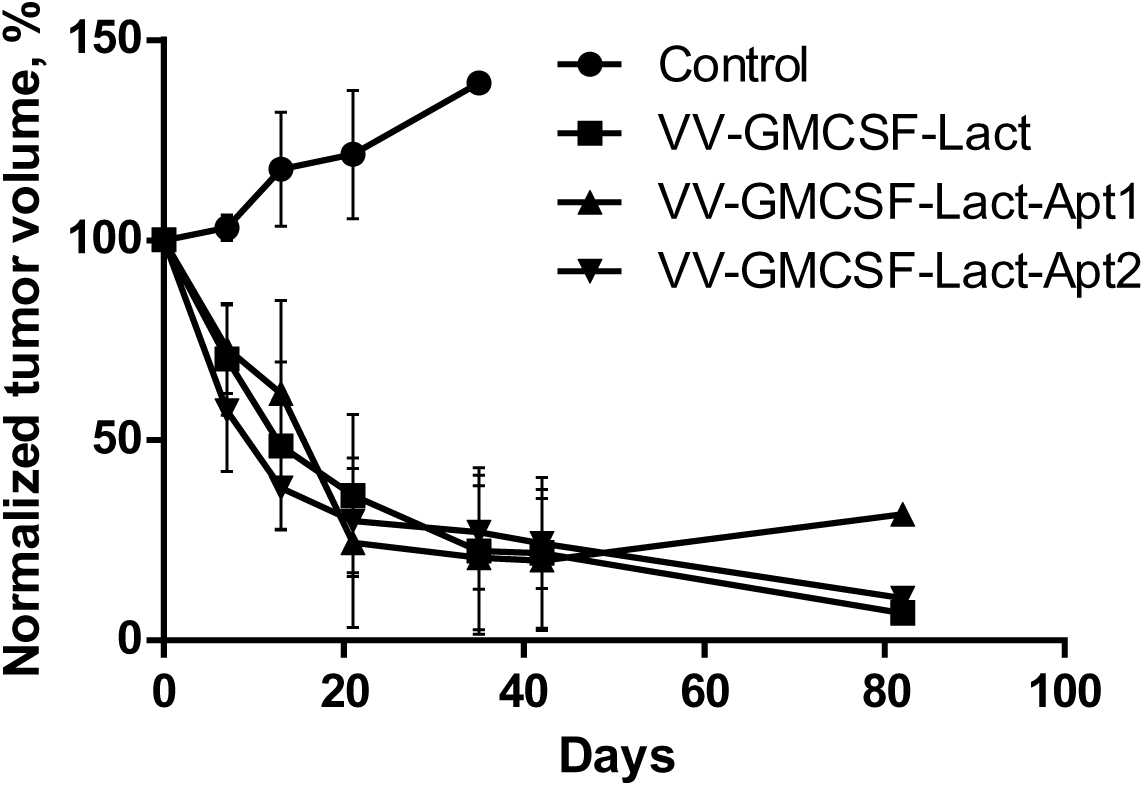
Tumor growth dynamics in response to a 4-week course of therapy with viral preparations (VV-GMCSFLact-Apt1, VV-GMCSF-Lact-Apt2 and VV-GMCSF-Lact).

### 2. Analysis of survival of mice with orthotopically transplanted glial tumors after systemic treatment with viral preparations

We evaluated the survival of immunocompromised mice bearing orthotopic glioma xenografts after treatment with viral preparations (VV-GMCSFLact-Apt1, VV-GMCSF-Lact-Apt2, or VV-GMCSF-Lact). By the end of the three-month observation period, counted from the last saline injection, one in ten mice in the control group remained alive (survival rate 10%). When comparing the control and experimental groups, we obtained the following results (Figure 7). Using both the Gehan-Breslow-Wilcoxon test and the log-rank test (Mantel-Cox test), statistically significant differences were found between the VV-GMCSF-Lact-Apt1, VV-GMCSF-Lact-Apt1 and control groups (p-value < 0.01). Treatment with the oncolytic virus VV-GMCSF-Lact alone resulted in a three-month survival rate of 36.4% (4 of 11 animals). Survival strongly correlated with the animals’ clinical condition. The four surviving mice displayed no visible abnormalities, maintaining a normal phenotype and body weight within the reference range. However, the majority of animals in this group (6 of 11) developed adverse effects, primarily manifested as deterioration in coat quality and significant weight loss.

**Figure 7.**
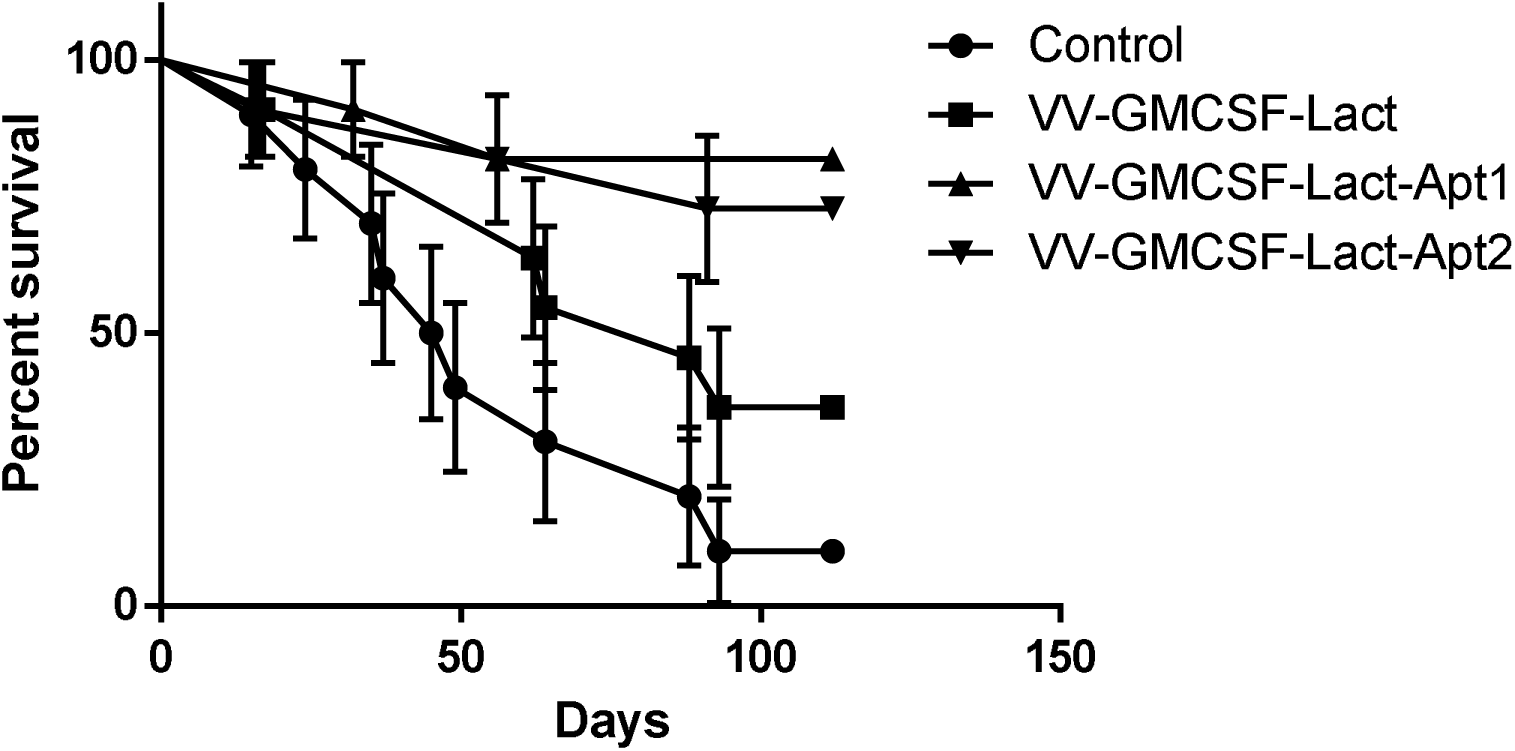
Survival curves of mice following a course of treatment with VV-GMCSFLact-Apt1, VV-GMCSF-Lact-Apt2 and VV-GMCSF-Lact.

In the VV-GMCSFLact-Apt1 group, a course of treatment induced a complex of negative clinical signs, including severe hypodynamia, deterioration in coat quality (sparse, dull, and ruffled fur), and significant weight loss. Despite these pronounced signs of toxicity, the three-month survival rate was 81.8% (9 of 11 animals). Of the two fatalities, one was directly attributed to intergroup aggression (cannibalism), precluding its interpretation as a treatment-related death.

Conversely, in the VV-GMCSFLact-Apt2 group, no adverse clinical signs were observed following the treatment course. This group achieved a three-month survival rate of 72.7% (8 of 11 animals). Among the three deaths, one was also due to intergroup aggression.

### 3. Evaluation of the level of key cytokines

To assess the level of key cytokines (IL-1, IL-10, TNF-alpha) in plasma, mice bearing xenografts were injected intravenously with the viral preparations (VV-GMCSF-Lact-Apt1, VV-GMCSF-Lact-Apt2 and VV-GMCSF-Lact). Cytokine levels were determined by ELISA after administration of the viral preparations at 1th day and 3th day and on the 5th day (Figure 8).

**Figure 8.**
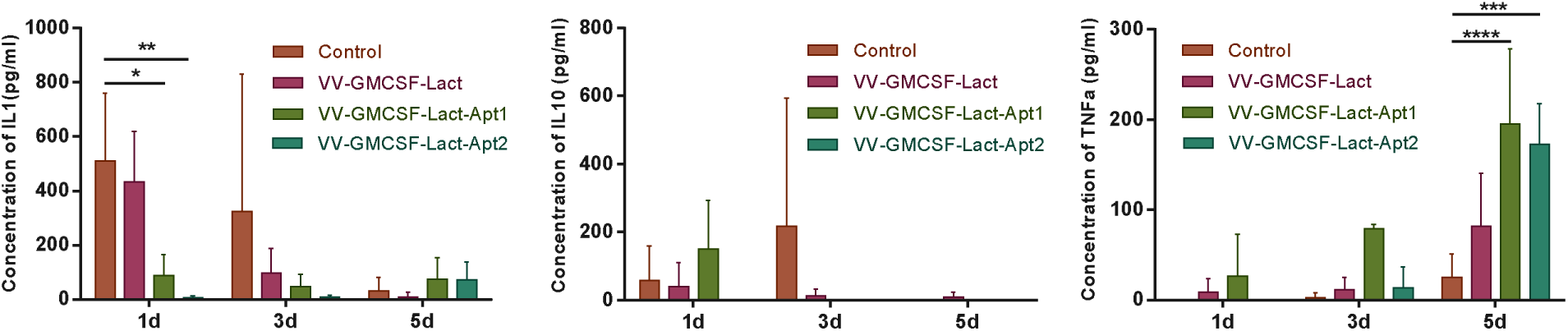
Levels of key cytokines (IL-1, IL-10, TNF-alpha) in the plasma of mice after administration of viral preparations VV-GMCSFLact-Apt1, VV-GMCSF-Lact-Apt2 and VV-GMCSF-Lact.

When comparing the mean values of IL1 cytokine concentrations in the experimental groups, we found statistically significant differences only at the 24-hour point (1th day). The IL1 concentration was significantly lower in the group of animals administered viral preparations containing aptamers: VV-GMCSF-Lact-Apt1 (p<0.05), VV-GMCSF-Lact-Apt2 (p<0.01) compared to the control group. Overall, a decrease in IL1 levels was observed in the analyzed time range. A sharp increase in the level of anti-inflammatory IL10 in the control group was shown on the third day of drug administration, but on the fifth day there were only trace amounts. The level of TNFα increased throughout the entire observed period, and especially significantly by day 5 in the groups of animals with administration of viral preparations (VV-GMCSF-Lact-Apt1 (±195.2 vs ±24.9, p<0.001), VV-GMCSF-Lact-Apt2 (±172.4 vs ±24.9, p<0.005) compared to the control.

### 4. Histological analysis

Histological analysis revealed that tumors in all PBS-treated control animals corresponded to malignant glioblastoma, characterized by cellular polymorphism, nuclear atypia, and active proliferation (Figure 9A). Atypical tumor cells of medium and small size with deformed, hyperchromatic nuclei and markedly scant cytoplasm (a sign of pronounced anaplasia and low differentiation). Occasional scattered lymphoid cells are present.

**Figure 9.**
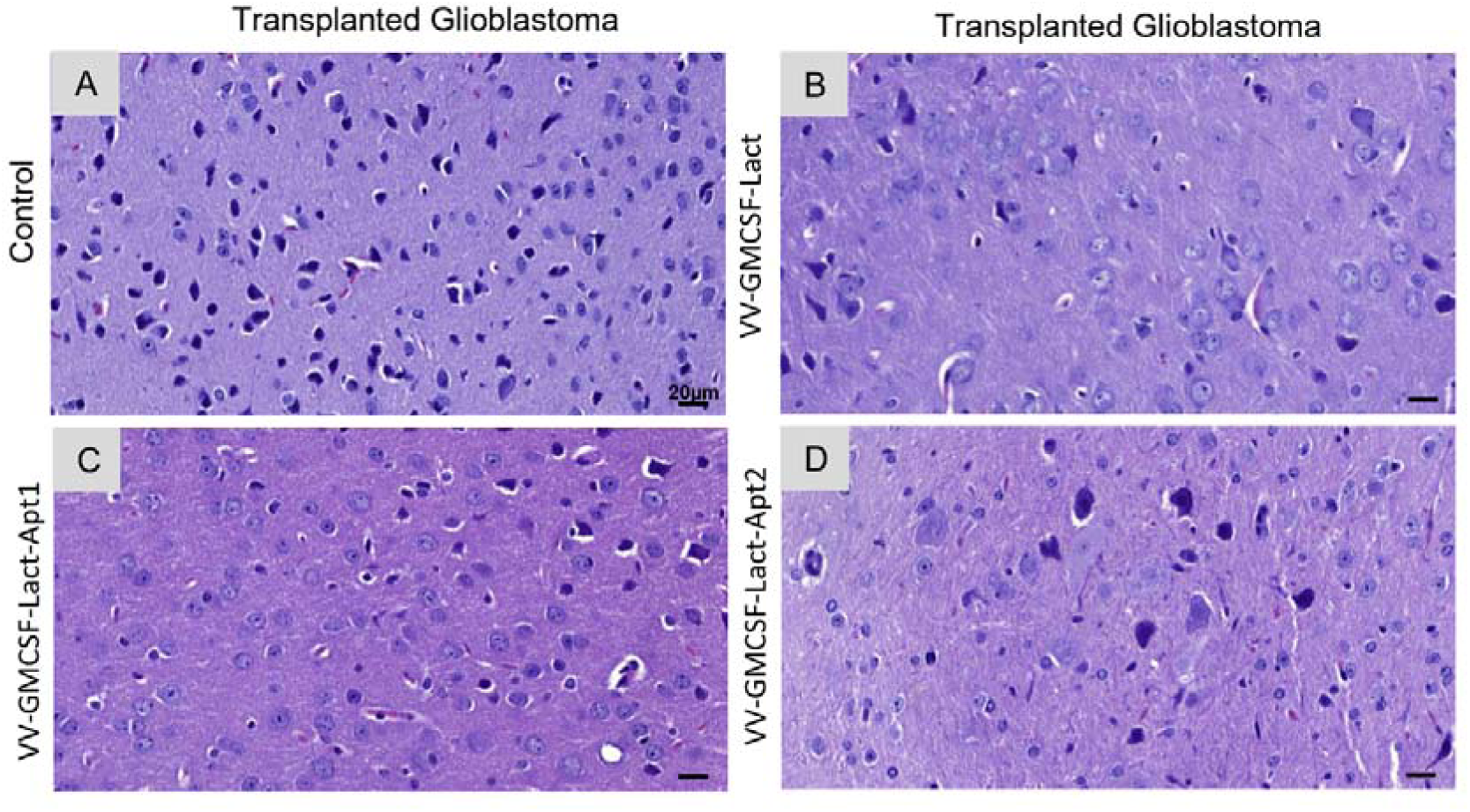
Histopathological analysis of glioblastoma xenografts after oncolytic virotherapy. Shown are representative hematoxylin and eosin (H&E)-stained sections of brain tumors from the following groups: (A) Control, (B) VV-GMCSF-Lact, (C) VV-GMCSF-Lact-Apt1, and (D) VV-GMCSF-Lact-Apt2.

All animals in VV-GMCSF-Lact group (Figure 9B). show significant pathological changes, indicative of a direct cytopathic effect of the virus and the development of an anti-tumor and immune response. The cells are in a state of acute viral cytopathic stress, which inevitably leads to their death (confirmed by the presence of “ghost cells”). There are small areas in the brain tissue that appear spongy and less dense, indicating possible edema (swelling) or necrosis. The reduction in viable cells ranges from approximately 20% to 30%, varying between animals and across different areas of the tumor. Moderate, focal lymphoid infiltration is noted, evidencing the initial activation of an immune response. Occasional cells show signs of apoptosis (pyknosis).

Compared to monotherapy with the virus, an increase and complication of pathological processes is observed in the VV-GMCSF-Lact-Apt1 group (Figure 9C). Alongside “ghost cells,” the number of cells with condensed and fragmented chromatin (pyknosis, karyorrhexis) increases, indicating the activation of apoptotic and necroptotic pathways. The plasma membrane of some cells becomes uneven. There are small areas in the brain tissue that appear spongy and less dense, indicating possible edema (swelling) or necrosis. The reduction in viable cells ranges from approximately 20% to 30%, varying between animals and across different areas of the tumor. Aptamer 1 enhances the efficacy of viral therapy, leading to more extensive infection and more diverse (mixed) tumor cell death, which morphologically corresponds to a more powerful pro-inflammatory stimulus. This may represent a later stage of infection, or perhaps the aptamer allowed the virus to reach the tumor more effectively and earlier.

VV-GMCSF-Lact-Apt2 group (Figure 9D) demonstrates the most pronounced pathomorphological changes. There is a predominance of “ghost cells” and large cellular debris with ragged edges and karyolysis. Cells that retain their structure are large (marked oncosis), with multiple nucleoli. A specific sign appears – a perinuclear halo (clearing around the nucleus), indicating severe damage to the endoplasmic reticulum, Golgi apparatus and an active viral cytopathic effect. Foci of cells with pyknosis and karyorrhexis persist. There are small areas in the brain tissue that appear spongy and less dense, indicating possible edema (swelling) or necrosis. The reduction in viable cells ranges from approximately 20% to 40%, varying between animals and across different areas of the tumor. An immune infiltrate is present, foci of lymphocytes and macrophages in the tumor tissue.

Histological analysis of organ tissue revealed only background drug toxicity caused by immunosuppression with cyclophosphamide, cyclosporine, and ketoconazole. No morphological differences were detected in the internal organ cells of mice from different groups. This confirms the selectivity of the oncolytic virus and the absence of additional organ-specific toxicity, reinforcing the arguments in favor of its favorable safety profile.

## DISCUSSION

Oncolytic viruses are a promising antitumor drug because they infect and replicate tumor cells, indirectly activating antitumor immunity ^6^. Their safety and efficacy have been confirmed in numerous preclinical and clinical trials. These viruses can be used either as monotherapy or in combination with immunotherapy, chemotherapy, or radiation therapy. It is worth noting that the clinical trials of VV-GMCSF-Lact (NCT05376527) utilized intratumoral administration of the oncolytic virus. Alternative routes of virus administration include intrathecal administration to achieve maximum drug concentrations in brain tissue and cerebrospinal fluid, as well as technically easier intravenous administration. However, the presence of neutralizing antibodies in patients may reduce antitumor efficacy with this route of administration ^7^. Aptamers, which can shield the oncolytic virus from neutralizing antibodies, may be a solution to this problem ^8^. Aptamers can also prevent the aggregation of viral particles and act as cryoprotectants during transportation, which increases their antitumor effectiveness ^9, 10^.

We previously identified aptamers that efficiently bound VV-GMCSF-Lact virus, were stable in the presence of serum nucleases, and prevented viral particle aggregation ^5,11^. Some of these aptamers (Nv14t_56 (№9), NV1t_72 (№1), NV4t_64 (№4), NV4t_53 (№5), NV14t_41 (№8)) were also shown to have a synergistic effect in vitro experiments on immortalized and primary glioblastoma cell lines. The aim of this study was the prove of the principle - whether aptamers in combination with viruses (VV-GMCSF-Lact-Apt1, VV-GMCSF-Lact-Apt2) would exert a synergistic antitumor effect in an orthotopic glioblastoma xenograft model.

In the in vivo experiment, we used ICR mice with incomplete immunosuppression as an animal model ^12^. This drug-induced immunosuppression mouse model, on the one hand, allows for the grafting and production of human glioblastoma xenografts, and on the other hand, when discontinuing the drugs (cyclosporine, cyclophosphamide, ketaconazole), it is possible to obtain a certain immune response when administering the vaccinia virus. Analyzing the MRI images, we see that tumor size decreased in all experimental groups and was not statistically different from each other (Figure 6). Perhaps we should have continued scanning until the end of the experiment—on day 112—to see if there were differences between the viral drug groups. We showed that mice treated with viral preparations containing aptamers (VV-GMCSF-Lact-Apt1, VV-GMCSF-Lact-Apt2) survived longer than mice treated with the virus alone (VV-GMCSF-Lact), not to mention control mice, whose survival was low (Figure 7). It is worth noting that despite the better survival, the group of mice administered with the VV-GMCSF-Lact-Apt1 were characterized by a complex of negative clinical signs, including severe hypodynamia, deterioration of coat quality (sparse, dull, and ruffled fur), and significant weight loss.

Here we used animals with incomplete immunosuppression, in which the effect lasts for approximately 14 days. But it is worth noting that even when using high doses of cyclophosphamide 150 mg/kg, the maximum changes in the parameters of the cellular and humoral immunity of animals were recorded on the 5th day of the study with a gradual recovery by the 18th ^12^. The presence of neutralizing antibodies after the first administration of the vaccinia virus was demonstrated in volunteers, providing a pronounced specific humoral and T-cell immune response ^13^. It can be assumed that by the third or fourth injection, neutralizing antibodies could have already been developed, but the presence of antiviral drugs protects viruses from their effects.

In this work, we investigated the level of first-generation proinflammatory cytokines (IL-1β, TNF-alpha), which initiate the inflammatory response, and the level of anti-inflammatory cytokines (IL-10), which regulate specific immune responses and limit the development of inflammation ^14^. Activated antigen-presenting cells synthesize IL-1β and TNF-alpha, resulting in the activation and further differentiation of T and B lymphocytes, increased expression of acute phase proteins, enhanced phagocytosis, and activation of endothelial cells. However, IL-1β also influences the tumor microenvironment, promoting its initiation and progression, while TNF-alpha suppresses tumor cell proliferation and induces tumor regression ^15^. The anti-inflammatory cytokine IL-10 is secreted by T helper type-2 cells, which in turn promote antibody synthesis by B cells (the productive phase of the humoral immune response)^16^. This process results in the formation of effector plasma cells, which begin producing antibodies of only one specificity. A study of the level of cytokines (IL-1, IL-10 and TNF-a) showed several patterns: starting from the first day, the level of IL-1 decreases, especially in the groups (VV-GMCSF-Lact-Apt1, VV-GMCSF-Lact-Apt2), and by the 5th day, the level of TNF-a begins to increase, which may indicate a shift in the immune response from a pro-inflammatory systemic one, low IL1, towards an anti-tumor one - an increase in TNF.

Thus, we demonstrated that viral preparations containing aptamers are more effective against orthotopic xenografts. This represents the first proof of principle that aptamers have a synergistic effect in combination with the VV-GMCSF-Lact virus in an *in vivo* experiment.

The antitumor efficacy of the viral preparations VV-GMCSFLact-Apt1, VV-GMCSF-Lact-Apt2, and VV-GMCSF-Lact against glioblastoma xenografts obtained by orthotopic transplantation of primary 3D glioma cultures into mice with incomplete drug-induced immunosuppression was demonstrated. No significant differences in tumor volume reduction were observed between the experimental groups. The survival rate of glioblastoma-bearing mice treated with VV-GMCSF-Lact-Apt1 and VV-GMCSF-Lact-Apt2 was higher than that of mice treated with the oncolytic drug alone (81,8% and 72,7% vs 36,4%, respectively). Thus, aptamers exhibit a synergistic effect when combined with an oncolytic virus, nearly doubling the survival of mice with glioblastoma. A study of cytokine levels (IL-1, IL-10, and TNF-α) showed that the aptamers shift the immune response from a pro-inflammatory systemic response, by reducing IL-1, to an anti-tumor one, with an increase in TNF-α.

Histological analysis revealed the most destructive changes in tumor tissue in the VV-GMCSF-Lact-Apt2 group - the scale and diversity of tumor cell death progress from isolated lysis to more massive death, involving lytic, apoptotic, and necroptotic mechanisms. Thus, based on the cytotoxicity, antitumor activity, and survival studies in animal models, it can be concluded that the viral preparations GMCSF-Lact-Apt1 and VV-GMCSF-Lact-Apt2 are more effective against orthotopic human glioblastoma xenografts than V-GMCSF-Lact, suggesting a synergistic effect of aptamers in combination with VV-GMCSF-Lact.

## MATERIALS AND METHODS

PBS, Fetal bovine serum FBS (Dia-M, Russia); Sodium bicarbonate (Sigma-aldrich, USA); Antibiotic Antimycotic Solution 100× liquid, Gibco® MEM Non-Essential Amino Acids, DMEM/F12, GlutaMAX, TrypLE, Dulbecco’s phosphate-salt buffer (Gibco, USA); Trypan blue, Tween20 (Bio-Rad Laboratories, USA); cyclosporine, cyclophosphamide, and ketoconazole, Zoletil (Virbac SA, France); Romethar (Interchemie, The Netherlands); аptamers to VV-GMCSF-Lact were synthesized by Lumiprobe, Russia.

### 1. Glioma cell cultivation

Culture of primary tumor cells from human glioblastomas were obtained by mechanical and enzymatic destruction of glioblastoma (G4 according to WHO), resected during surgery in a patient in 2022. The material was collected with the approval of the Local Ethics Committee (LEC) of the Federal State Budgetary Educational Institution of Higher Education “Krasnoyarsk State Medical University named after Professor V.F. Voyno-Yasenetsky” of the Ministry of Health of the Russian Federation (Protocol No. 95/2020 dated 01/29/2020), as well as with the approval of the bioethics commission of the LEC of KrasSMU dated 11/05/19 and the LEC of the “Krasnoyarsk Interdistrict Clinical Emergency Hospital named after N.S. Karpovich” dated 11/20/2016. Culture of glioblastoma cells was cultured in DMEM/F12 with the addition of 10% FBS, 1X solutions of GlutaMAX, 1X solutions of Gibco® MEM Non-Essential Amino Acids and antibiotic-antimycotic at 37°C in a CO2 incubator.

### 2. Oncolytic Virus

Recombinant VV-GMCSF-Lact was engineered from L-IVP vaccinia virus and has deletions of the viral thymidine kinase (tk) and vaccinia growth factor (vgf) gene fragments, in the regions of which the genes of human GM-CSF and the oncotoxic protein lactaptin are inserted ^3^.

### 3. Formation of aptamer complexes with the virus

VV-GMCSF-Lact virus at a concentration of 3.6×107 PFU/ml was incubated for 30 minutes at 25 C with 50 µl of 200 nM pre-annealed aptamers Nv14t_56 (№9) – the viral preparation VV-GMCSF-Lact-Apt1; with a pool of aptamers (NV1t_72 (№1), NV4t_64 (№4), NV4t_53 (№5), NV14t_41 (№8)) – the viral preparation VV-GMCSF-Lact-Apt2.

### 4. Experimental animals

The study was carried out in compliance with the ARRIVE guidelines. All experiments with laboratory animals were carried out in accordance with the recommendations and requirements of the World Society for the Protection of Animals (WSPA) and the European Convention for the Protection of Vertebrate Animals (Strasbourg, 1986). Permission for experiments with laboratory animals was obtained from the Bioethics Committee for Work with Animals of the Local Ethics Committee of the State Budgetary Educational Institution of Higher Education Krasnoyarsk State Medical University named after prof. V.F. Voyno-Yasenetsky of the Ministry of Health of the Russian Federation on December 16, 2022, Extract from Protocol No. 3. In vivo experiments were conducted using female ICR laboratory mice weighing 20-25 g, housed in sterile, individually ventilated cages. Animal housing, feeding, care, and withdrawal from the experiment were carried out in accordance with the requirements of the “European Convention for the Protection of Vertebrate Animals used for Experimental and Other Scientific Purposes” (Strasbourg, 1986) and the “Rules for Conducting Work Using Experimental Animals” (Order 755 of August 12, 1977, USSR Ministry of Health). Mice were housed 5 per cage. They were examined daily, and their general condition was assessed. Before treatment, mice were divided into groups of 10 animals, depending on the treatment they were receiving. Animals that were allowed to survive according to the experimental protocol continued to be examined daily, and their general condition was assessed, for three months after the experiment. After three months (five-year survival), the mice were removed from the experiment.

### 5. Obtaining mice with incomplete drug-induced immunosuppression

Partial drug immunosuppression was performed using the following drugs, according to the following regimen: cyclosporine (20 mg/kg intraperitoneally), cyclophosphamide (60 mg/kg intraperitoneally), and ketoconazole (10 mg/kg orally) every second day for 10 days before and 2 days after tumor transplantation.

### 6. Orthotopic transplantation of glioblastoma cells into immunosuppressed animals

For establishing brain tumor orthotopic xenografts, immunodeficient mice, 4- to 5-week-old mice were anaesthetised with Zoletil (40 mg/kg, 50 μL) and Romethar (50 μL) and fixed through the ear canals in a stereotaxic system (RWD Life Science, China). Mice were scalped and a trepanation hole was made in the skull in the projection of the striatum of the right cerebral hemisphere using Drill Bits HM1005 0.5 mm, Round Tip (RWD Life Science, China). Culture of primary tumor cells (100,000 cells in 5 μL of sterile Dulbecco’s phosphate-salt buffer) were injected using a micropump at a rate of 1 μL/min into the brain striatum: A-1+ D1,5 V-2,5mm.

### 7. Monitoring Brain Tumor Development Using Omniscan Contrast-Enhanced MRI

Imaging and Experimental Timeline. Tumor growth was monitored using contrast-enhanced Magnetic Resonance Imaging (MRI) with Omniscan (0.5 mmol/ml, GE Healthcare Ireland Limited). The initial MRI scan was performed on day 14 after glial cell transplantation, prior to the initiation of therapy with the viral agents VV-GMCSF-Lact-Apt1, VV-GMCSF-Lact-Apt2, and VV-GMCSF-Lact. For the single-treatment regimen, subsequent imaging was performed on day 7 after viral administration. For the continuous-treatment regimen, MRI was performed every 7 days for 28 days, with a final scan 2 weeks after the fourth drug administration.

Assessment of Therapeutic Efficacy. The efficacy of single and repeated intravenous administrations of the viral preparations was assessed by comparing brain MRI scans from mice with transplanted tumors. Comparisons were made between scans taken before therapy, after a single administration, and every 7 days throughout the continuous treatment course.

MRI Acquisition Parameters. MRI was performed on mice under anesthesia, following a modified protocol established in a previous study ^17^. Scans were conducted using an Avance DPX 200 spectrometer (Bruker BioSpin GmbH) with the contrast agent Omniscan. Two-dimensional, slice-selective images were acquired using a multi-slice multi-echo sequence (Paravision 4.0 software, Bruker BioSpin). The acquisition parameters were as follows: Slice Thickness: 0.71 mm; Field of View: 40 mm; Matrix Size: 256 × 256 pixels; In-Plane Spatial Resolution: 156 µm/pixel; Repetition Time (TR): 600 ms; Echo Time (TE): 4.7 ms (to generate T1-weighted contrast); Total Acquisition Time: 10 minutes.

Tumor Volume Quantification. Tumor volume (in mm³) was calculated from the MRI sections according to the following protocol:

The tumor area in pixel² was measured on all slices where the tumor was present. The area in pixel² was converted to mm² using a 1 cm reference scale present on each MRI image. The conversion factor (denoted as S in the formula) varied between experimental series depending on image post-processing and conversion to JPEG format. Each resulting cross-sectional area (in mm²) was multiplied by the slice thickness (0.71 mm) to obtain the volume for that individual slice. The volumes of all slices were summed to obtain the total tumor volume. The formula for tumor volume calculation is:

Tumor Volume = Σ (Area_slice_n [pixel²]) / S × 0.71 mm, where *n* is the number of slices containing the tumor focus, and S is the conversion factor (pixels² to mm²) determined from the scale on the image.

The normalized tumor volume was calculated as follows:

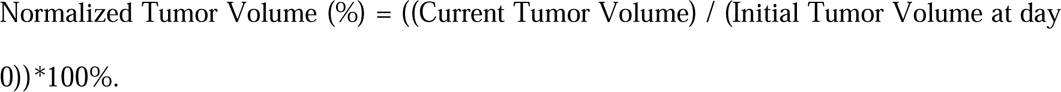

### 8. The enzyme linked immunosorbent assay

Blood plasma was collected from the animals during euthanasia at 24 hours, 72 hours, and 5 days after the initiation of therapy. For each time point, plasma samples were obtained from 3 animals (n=3 per group). The enzyme linked immunosorbent assay for cytokines IL-1β, IL-10, TNF-α was performed using commercial kits (catalog number E-EL-M0037, E-EL-M0046, E-EL-M3063, respectively, Elabscience, USA) according to the manufacturer’s instructions.

### 9. Histological analysis

For the histological study, the brain specimens with tumor nodes were fixed in 10% neutral-buffered formalin (BioVitrum, Moscow, Russia), dehydrated in ascending ethanols and xylols and embedded in HISTOMIX paraffin (BioVitrum, Russia). The paraffin sections (up to 5 µm) were sliced on a Microm HM 355S microtome (Thermo Fisher Scientific, Waltham, MA, USA) and stained with hematoxylin and eosin. The images were examined and scanned using an Axiostar Plus microscope equipped with an Axiocam MRc5 digital camera (Zeiss, Oberkochen, Germany) at magnification of ×100, ×200 and ×400.

### 10. Statistical analysis

Statistical analysis was performed using GraphPad 6.01 (GraphPad Software, San Diego, CA, USA). Quantitative variables are presented as mean ± standard deviation (SD). A two-way ANOVA was used to compare more than two data sets. The survival data were analyzed by Kaplan-Meier survival analysis. The Gehan-Breslow-Wilcoxon test and the log-rank test (Mantel-Cox test), was used to compare two samples or groups. Differences were considered significant if the P-value was <0.05.

## Supplementary Materials

The following supporting information can be downloaded at: https://www.mdpi.com/article/doi/s1, Figure S1: title; Table S1: title; Video S1: title.

## Author Contributions

Conceptualization, M.D., A.K., E.K.; methodology, M.D., A.A., N.V., A.K., N.L.; software, M.D., X.X.; validation, M.D., A.A., N.V., A.K., N.L.; formal analysis, M.D.; investigation, A.K., E.M., V.K., A.K., N.L., T.G., O.S. resources, A.S., V.R.; data curation, M.D., N.L.; writing—original draft preparation, M.D., A.K., N.L., E.M., N.V., O.S.; writing—review and editing, M.D., A.K., E.M., V.K.; visualization, A.K.; supervision, A.K., V.R.; project administration, V.R.; funding acquisition, E.K. All authors have read and agreed to the published version of the manuscript.

## Institutional Review Board Statement

The material was collected with the approval of the Local Ethics Committee (LEC) of the Federal State Budgetary Educational Institution of Higher Education “Krasnoyarsk State Medical University named after Professor V.F. Voyno-Yasenetsky” of the Ministry of Health of the Russian Federation (Protocol No. 95/2020 dated 01/29/2020), as well as with the approval of the bioethics commission of the LEC of KrasSMU dated 11/05/19 and the LEC of the “Krasnoyarsk Interdistrict Clinical Emergency Hospital named after N.S. Karpovich” dated 11/20/2016. Animal housing, feeding, care, and withdrawal from the experiment were carried out in accordance with the requirements of the “European Convention for the Protection of Vertebrate Animals used for Experimental and Other Scientific Purposes” (Strasbourg, 1986) and the “Rules for Conducting Work Using Experimental Animals” (Order 755 of August 12, 1977, USSR Ministry of Health).

## Informed Consent Statement

Informed consent was obtained from all subjects involved in the study.

## Data Availability Statement

Data are contained within the article.

